# Mesopredator-mediated trophic cascade can break persistent phytoplankton blooms in coastal waters

**DOI:** 10.1101/2022.06.07.495132

**Authors:** Maximilian Berthold, Rhena Schumann, Volker Reiff, Rita Wulff, Hendrik Schubert

## Abstract

Managing eutrophied systems only bottom-up (nutrient decreases) can be economically and ecologically challenging. Top-down controls (consumption) were sometimes found to effectively control phytoplankton blooms. However, mechanistic insights, especially on possible trophic cascades, are less understood in brackish, species-poor coastal waters, where large cladocera are absent. In this study, we set-up large mesocosms for three consecutive years during growth season. One set of mesocosms was controlled by mesopredator (gobies and shrimp), whereas the other mesocosms had no such mesopredator present. The results were standardized to monitoring data of the ecosystem to denote possible differences between treatments and the system. We found that mesopredator mesocosms showed lower turbidity, phytoplankton biomass, and nutrients compared to no-mesopredator mesocosms and the ecosystem. This decrease allowed macrophytes to colonize water depths only sparsely colonized in the ecosystem. Rotifer biomass increased in mesopredator mesocosms compared to the ecosystem and no-mesopredator mesocosms. Likewise, copepod biomass that potentially grazes upon rotifers and other microzooplankton decreased in mesopredator mesocosms. No-mesopredator mesocosms were colonized by an omnivorous mesograzer (*Gammarus tigrinus*), potentially creating additional pressure on macrophytes and increasing grazing-mediated nutrient release. Zooplankton was not able to control the non-nutrient limited phytoplankton. We propose a new mechanism, where a higher mesopredator density will increase grazing on phytoplankton by promoting microzooplankton capable of grazing on picophytoplankton. This proposed mechanism would contrast with freshwater systems, where a decrease of zooplanktivorous fish would promote larger phytoplankton grazer like cladocera. Biomanipulation in such species-poor eutrophic coastal waters may be more successful, due to less trophic pathways that can cause complex top-down controls. Stocking eutrophic coastal waters with gobies and shrimps may be an alternative biomanipulative approach rather than selectively remove large piscivorous or omnivorous fish from eutrophic coastal waters.

## Introduction

One of the biggest challenges in ecosystem restoration is restoring deteriorated aquatic ecosystems back to an undisturbed state in the coming decades. One factor that still affects ecosystems until nowadays is eutrophication. The main approaches developed to control eutrophication have been to reduce the influx of elements, like Nitrogen, and Phosphorus (Schindler 1977, Vitousek et al. 1997, Schindler et al. 2016). These decreases worked mostly for deep lakes, where pre-eutrophication Chlorophyll concentrations were reached after three decades, but with different plankton species compositions than before eutrophication (Jochimsen et al. 2013). Larger bodies of water, like the Baltic Sea would not reach pre-disturbed phosphate concentrations within the next 100 years, at the current rate of nutrient concentration decrease, due to insufficient inflow decrease in some areas (HELCOM 2014). Shallow aquatic ecosystems are even more affected, as they lack the water depth for long-term nutrient sequestration. Lake Tegel, a shallow German lake reached pre-disturbed ecosystem states through large external measures, like diverting and treating inflowing waters, within a little more than 30 years (Chorus et al. 2020). Such approaches aimed at bottom-up control can become unfeasible, especially, if management practices do not improve within the surrounding catchment area. For example, still increasing phosphorus stocks in soils (van Dijk et al. 2016), can cause ongoing phytoplankton growth in shallow coastal waters through nutrient flushing (Berthold et al. 2018a). However, shallow lagoons are very important, globally, as they make up 10% of the global coastline (Dürr et al. 2011), and locally act as a nutrient filter for the Baltic Sea (Asmala et al. 2017).

Biomanipulation through top-down control has been proposed as an additional sustainable method to support restoration efforts (Jeppesen et al. 2007). Top-down control can include stocking of piscivorous fish (pike, sander), which in turn control zooplanktivorous fish, favoring phytoplankton consumption by zooplankton. There are a wide variety of top-down biomanipulative studies published, either conducted in mesocosms, or in whole lakes (Jeppesen et al. 1990, Søndergaard et al. 2007). Most studies have described an effective decrease of phytoplankton biomass, depending on the ratio of zooplanktivorous to piscivorous fish, and the amount of large cladocera as unselective filter feeders. The majority of experiments focused on shallow, limnic systems, but shallow coastal waters are similarly impacted by eutrophication. Such brackish systems show rarely, if any large cladocera, and as such grazing rates on phytoplankton can be low in certain lagoons of the Baltic Sea (0.2 - 6% of pelagic primary production, Heerkloss et al. 1984, Schiewer 2007). However, there are studies describing potential trophic cascades in coastal waters (Compte et al. 2012) and metabarcoding has revealed new trophic pathways between zooplankton-phytoplankton of various size classes in the Baltic Sea (Zamora-Terol et al. 2020, Novotny et al. 2021). Coastal waters are furthermore connected to limnic and marine systems and are hatching points for migratory fish species, like herring and perch (Aro 1989). A total removal of one fish species like in enclosed systems (lakes) is therefore not possible. Such prerequisites make biomanipulative top-down approaches in shallow coastal waters challenging. Nonetheless, those approaches need to be explored, if nutrient decreases by point sources cannot be lowered any further.

Furthermore, the Baltic Sea is already more impacted by increased sea surface temperature, compared to other areas in the world (Reusch et al. 2018). This temperature, and nutrient condition acts upon the functioning of the food web, by promoting a decrease in body size of every compartment of the food web (Peter and Sommer 2013, Liénart et al. 2021). If increasing temperatures decrease the cell size of phytoplankton (Sommer et al. 2017), than the food web would equally respond selecting smaller individuals (Liénart et al. 2021). Such small phytoplankton is problematic, as it may pass zooplankton grazing (Lürling 2020), therefore lowering food web efficiency. One such system with perennial dominance of picophytoplankton is the Darß-Zingst lagoon system (DZLS, Albrecht et al. 2017), which is located at the southern Baltic Sea coast of Germany. The DZLS is a typical lagoon system of the southern Baltic Sea and acts as a model ecosystem, due to its salinity and trophic gradient (salinity of 0 - 13 PSU, highly eutrophic - mesotrophic, Schiewer 2007). This system is 90% dominated by cyanobacteria, showing simultaneously a high TN:TP of 35:1, and no available dissolved inorganic P during summer (Berthold 2016, Berthold et al. 2018b, Schumann et al. 2019). There is a fluctuating dominance of copepods over rotifers throughout the year (Feike et al. 2007), and grazing of zooplankton is assumed to be only around 3% of the gross primary production (based on mean, Heerkloss et al. 1984). In theory, such low grazing rates would point to a missing top-down control within the ecosystem, where the amount of planktivorous fish is too high. However, there have been increasing amounts of reports, that such food web interactions are too simple. For example, grazing on copepods can act as switch within trophic cascades (Stibor et al. 2004). Strong grazing on copepods can support ciliates, which graze upon small phytoplankton, whereas weak grazing on copepods promotes their grazing on ciliates and large algae favoring small phytoplankton (Stibor et al. 2004). Smaller phytoplankton is in turn more efficient at lower nutrient concentrations shifting the overall plankton community composition (Schmidt et al. 2020). Such trophic cascades can change the composition of micro- and mesozooplankton communities (Zöllner et al., 2009), ultimately affecting a zooplankton-mediated top-down control on phyto- and bacterioplankton (Zamora-Terol et al. 2020). In the case of the DZLS, such a possible trophic cascade has been observed, with grazing of *Eurytemora affinis* on eggs and adults of the rotifer *Keratella cochlearis* (Feike and Heerkloss 2009). However, rotifers such as *K. cochlearis* are known to graze upon bacterioplankton (Haukka et al. 2006), but also on filamentous cyanobacteria (Novotny et al. 2021), which provide a large fraction of total biovolume in the DZLS (Albrecht et al. 2017, Berthold and Schumann 2020). The DZLS shows a food web with few trophic levels, a high redundancy (Paar et al. 2022), and a dominant energy- and matter flow within the microbial loop (Schiewer et al. 1990, Schiewer 2007). An increased grazing pressure by rotifers and ciliates may disrupt the microbial loop, by lowering the picoplankton compartments that otherwise turn over nutrient pools through enzymatic activity within hours (Berthold and Schumann 2020).

We therefore questioned if restoration measures focusing on nutrient decrease were not successful, because a trophic cascade prohibited an efficient grazing on phytoplankton, especially if a trophic cascade than promotes small phytoplankton, which is potentially grazing resistant and highly competitive in a low-nutrient environment. We hypothesized that the removal of large predators, such as piscivorous fish (trophic level 3) leads to increased abundance of mesopredator such as gobies and shrimp (trophic level 3). Thus, higher abundances of mesopredator can potentially increase the consumption of mesograzer such as herbi- and omnivorous amphipods and isopods (trophic level 2) and zooplankton (copepods, cladocera). This increased consumption would cause a disruption of the pelagic and benthic nutrient recycling, ultimately promoting a clear water state by favoring smaller zooplankton grazing on smaller phytoplankton. Contrarily, if mesopredator do not create this postulated trophic cascade, a complete exclusion of all mesopredator would lead to very high numbers of zooplankton, promoting a clear water state. The exclusion of mesopredator would be equal to an efficient grazing control by large piscivorous fish like pike. The ecosystem would be used as control for the annual development of all monitored variables (nutrients, phytoplankton, zooplankton) in mesocosms. The main contrasting hypotheses can be formulated as follows:

i. Mesopredator presence will disrupt the microbial loop in a eutrophic lagoon through an intra-zooplankton trophic cascade, causing a clear water state.
ii. Absent mesopredator will liberate zooplankton of grazing pressure, increasing grazing rates on phytoplankton, causing a clear water state.

We used large *ex-situ* mesocosms for this biomanipulation and replicated them three years in a row, to catch possible interannual effects. Monitoring of the ecosystem and the mesocosms allowed to follow the seasonal development of all pelagic compartments.

## Material and Methods

### Experimental Set-up

The mesocosms were set-up on the premises of the Biological Station Zingst near the DZLS on the southern Baltic Sea coast. A general experimental set-up can be found in Berthold (2018). Briefly, mesocosms were 2200-liter glass fiber containers (dimensions: internal height 92 cm, internal width at base 119 cm, internal length at base 182 cm) and were embedded in the ground to allow for passive temperature control by the surrounding soil. Mesocosms stood on an unobstructed field without interference of any structure that would impact wind, precipitation, or sunlight. We followed design suggestions made by Raffaelli and Moller (1999) regarding replicates, independence and ecosystem comparability (see following points). The mesocosms were started in 2015, and were repeated in 2016, and 2017 for a total of five replicates per treatment over these three years. Each year, mesocosms were filled with new habitat sediment (5 cm total height) and habitat water in late spring. Mesocosms with and without mesopredator were assigned randomly to avoid pseudo-replicates (Raffaelli and Moller 1999). Habitat water was pumped in unfiltered from the surface water of the near-by lagoon system (50 m away) once at the start of each years’ experiment. We did not add additional habitat water to compensate for evaporated water to avoid increasing salinities. Water levels were near constant with evaporation being almost equal to precipitation during that time. We used three internal water pumps (pump capacity – 300 L h^-1^, Co. Neptun) to imitate the polymictic characteristics of the DZLS and prevent stratification within the mesocosms water columns. The upper 5 – 10 cm of sediment was collected each year at a water depth of 50 – 100 cm from a nearby ecosystem site (Grabow, Bodstedter Bodden) with known macrophyte colonization to guarantee a macrophyte diaspore bank allowing possible macrophyte sprouting. The sediment was thus not sieved. Thus, infauna was present in the mesocosms but not quantified. The mesocosms ran for a whole growth season, starting in April, and ending in September of each year, to include the annual macrophyte growth season. After the end of each season, mesocosms were emptied and cleaned. All biomass (pelagic, benthic) was collected and quantified (see below). Mesocosms of the first year were equipped with two optodes at two depths (surface ∼5 cm, and close to bottom at 70 cm) to test stability of primary production and effects of sediment respiration. Fish and shrimp were collected with a beach seine right before each experimental start. The beach seine had an opening of 3.5 m and was pulled along 100 m in a water depth from 1.5 – 0.5 m. Some further methodological remarks can be found in the Supplement section under “Methodological remarks on a possible “bottle effect” within the mesocosms”.

Fish and shrimp were kept in aerated habitat water containers, and their length determined before setting in mesocosms. We used the gobiid species *Pomatoschistus microps* and *P. minutus*, as well as the shrimp species *Palaemon elegans*. These species are abundant within the DZLS, and graze as mesopredator on larger zooplankton and mesograzer (amphipods, isopods) (Gagnon et al., 2021; Pasquaud et al., 2010; Pihl et al., 2006). *P. microps/minutus* were 2 – 3 cm long at the start of each experiment, which is a normal length during spring (Zander and Hartwig 1982). We added 10 gobies (1 – 1.5 g fresh mass m^-2^) and 10 shrimps (0.5 – 1 g fresh mass m^-2^) per mesocosm each year. The added amount of fish is within the range for similar estuaries based on abundance (5 -20 individuals m^-2^, Ythan estuary, Healey 1971, Jaquet and Raffaelli 1989) or fresh mass (∼ 1 g fresh mass m^-2^, island of Sylt, Zander and Hartwig 1982). The added shrimp biomass is within the range of other coastal water bodies in the Baltic Sea (Jephson et al. 2008, Moksnes et al. 2008). For simplicity, mesocosms with fish and shrimp added are labelled as mesopredator mesocosms throughout, whereas mesocosms without added fish and shrimp are labelled as no-mesopredator mesocosms.

### Mesocosm monitoring

Mesocosms were sampled frequently (biweekly) for abiotic and biotic parameters from their surface. Abiotic parameters included turbidity (optical density at 720 nm, m^-1^), dissolved inorganic phosphorus (DIP, µmol L^-1^), and dissolved inorganic nitrogen (DIN, sum of nitrate, nitrite, ammonium, µmol L^-1^), total phosphorus (TP, µmol L^-1^), and total nitrogen (TN, µmol L^-1^), and oxygen saturation (%). DIN and DIP was analyzed according to Hansen and Koroleff (1999). TN and TP were analyzed with an adjusted protocol using a sub-boiling extraction with potassium persulphate (Berthold et al. 2015). Determination limits were 0.26 µmol L^-1^ for nitrate, 0.6 for ammonium, 0.05 µmol L^-1^ for DIP, 3 µmol L^-1^ for TN, and 0.22 µmol L^-1^ for TP.

Water samples were filtrated onto glass fiber filters to determine Chlorophyll *a* concentrations (Chl *a*). Chl *a* was determined with a cold EtOH extraction overnight (Strickland and Parsons 1972) and Chl content was calculated following the standard HELCOM procedure (HELCOM 2013). Phytoplankton samples were taken weekly in the ecosystem, and at the end of each experiment in the mesocosms. Samples were preserved in Lugol (5 drops per 20 mL) and in glutaraldehyde (1% final concentration glutaraldehyde per sample) for later species determination and cell count using light and epifluorescence microscopy. Samples were refrigerated in the dark. We used sedimentation chambers (volume 1 mL, depth 3 mm) and a laboratory light microscope (Euromex, magnification 256x, 16x ocular, 16x objective) to count phytoplankton colonies and eukaryotic phytoplankton. Smaller phytoplankton and bacterioplankton, especially picocyanobacteria (<2 µm) were counted by epifluorescence microscopy (Olympus BX51). Plankton sub-samples (0.5 – 1 mL) were filtered onto Irgalan-black stained polycarbonate Track-Etch membranes (Sartorius 0.2 µm pore size, 25 mm). Filters were stained with DAPI solution for 5 minutes (1 mL, 92 µmol DAPI L^-1^, pH = 7.6) and cells were subsequently counted under the epifluorescence microscope (green excitation, U-MWG2 filter cube, UPlan FL 100 NA 1.3 oil objective). Biovolume of chlorophytes and heterokontophytes were calculated based on published cell sizes according to HELCOM (Olenina et al. 2006, HELCOM 2013). Cell sizes for picophytoplankton were based on electron microscopy cell sizes (Schumann, unpublished data).

Zooplankton was sampled at least four times over the course of each experiment fixating 1 L of a water sample with formaldehyde (4 % final concentration). The 1 L of full sample sedimented 24 h, 900 ml of supernatant were removed, and the sample was taken up in the remaining 100 ml. These were analyzed in total or in parts of at least 10 ml, depending on zooplankton abundances. An inverse microscope or Kolkwitz chambers were used for counting. Abundance numbers of occurring species were transformed to biomass (fresh mass µg L^-1^) using average geometric equivalents of individual body volumes according to a list by Heerkloss et al. (1991). This list also accounts for different size stages in copepods based on their life stage.

Animals and plants biomasses were collected at the end of each replicate run in September. Animals were counted and weighed. All animals were euthanized according to the regulations of the University of Rostock and the experiments were supervised by the animal welfare officer. We found three notable exceptions in our mesocosms at the end of the experiment. Three individuals of the invasive crab *Rhithropanopeus harrisii* were found in a mesopredator mesocosm in 2015, and one invasive goby *Neogobius melanostomus* was found in one mesopredator mesocosm in 2016. No-mesopredator mesocosms were colonized by gammarids (mostly *Gammarus tigrinus*) throughout all years (see Results, Discussion and Supplement for more information). Macrophyte biomass was harvested by hand, determined on genus level, blotted dry and wet-weighed. Biomass was then dried at 90 °C for 24 hours and re-weighed. Water content (%) was calculated based on differences between wet and dry mass. Fresh mass is reported throughout, but we used water content to confirm that fresh masses stated here are within the range of literature values and our own field sampling.

We further used the experimental and monitoring data to model seasonal dynamics of dissolved, and total nutrients, Chl *a*, turbidity, and zooplankton species. The monitoring station is situated at the Zingster Strom (54°25’47.0’’ N 12°41’14.5’’ E), right in the middle of the DZLS. Samples at this station were taken every day throughout the year for more than 40 years. Prior to modelling, we log-transformed and standardized our data. We chose to log-transform our data to make assumptions about relative changes within each year and treatment. Furthermore, we standardized our experimental data with the respective ecosystem monitoring data (daily measurements from Julian day 120 to 270, years 2015 – 2017) to make our data directly comparable to ecosystem trends (see Supplement Figure 1 for ecosystem averages of years 2015 – 2017). Standardization was carried out with the *scale* function of R (R Core Team 2018), which calculates a z-score of the respective data set. Zooplankton monitoring data was only available until December 2013 (see Supplement Figure 2 for ecosystem average per zooplankton group) and was only sampled once per week. We therefore used data from the last three available years (2011 – 2013) for standardization with our mesocosm data (2015 – 2017). We chose a generalized additive mixed model (GAMM, *gamm* function, R package *“mcgv”*, Wood 2011, Gaussian distributional assumptions, variable knots depending on data availability, k = 4 – 20, sampling year as random factor), as the best model to reflect the experimental data. A GAMM allowed us to include the sampling year as random factor in our analyses. We used the GAMM to predict the seasonal development as function of sampling date in both mesocosm experiments and the ecosystem. Furthermore, we were able to compare GAMM smooths for different treatments by including an ordered factor (treatment – ecosystem, mesopredator mesocosms, non-mesopredator mesocosms) to our models. We set the ecosystem data as reference level and calculated the difference smooths between reference level smoother (ecosystem) and treatment levels (mesocosms). We were thus able to describe statistically if the smooth terms per treatment and variable were significantly different from each other (Simpson 2017), that means ecosystem against mesocosms and the mesocosms treatments to each other.

## Results

### Pelagic elemental composition

Dissolved nitrogen (DIN) concentrations within mesocosms tended to increase above average in both treatments until June and would decrease below average in September (Figure 1A). Average values based on median can be found in Table 1. Concentrations within the ecosystem were rather stable throughout this period, with only a small decrease during August. The DIN smooths of both treatments were significantly different to the ecosystem for mesopredator (p < 0.0001) and no-mesopredator mesocosms (p < 0.0001). However, both treatments did not differ significantly to each other (p = 0.6). This development points to a higher DIN availability in both mesocosm treatments compared to the ecosystem. On the contrary, DIP concentrations in both mesocosm treatments, dropped during the month of June, with mesopredator mesocosms showing even lower concentrations compared to no-mesopredator mesocosms (Figure 1B). Contrary to the mesocosms, DIP concentrations within the ecosystem did not show any trends throughout the observational period. The DIP smooths of both treatments were significantly different to the ecosystem for mesopredator (p < 0.001) and no-mesopredator mesocosms (p = 0.024). However, both treatments did not differ significantly to each other (p = 0.64). Nonetheless, DIP concentrations were mostly close to the determination limit, pointing to overall low DIP concentrations.

**Figure 1.**
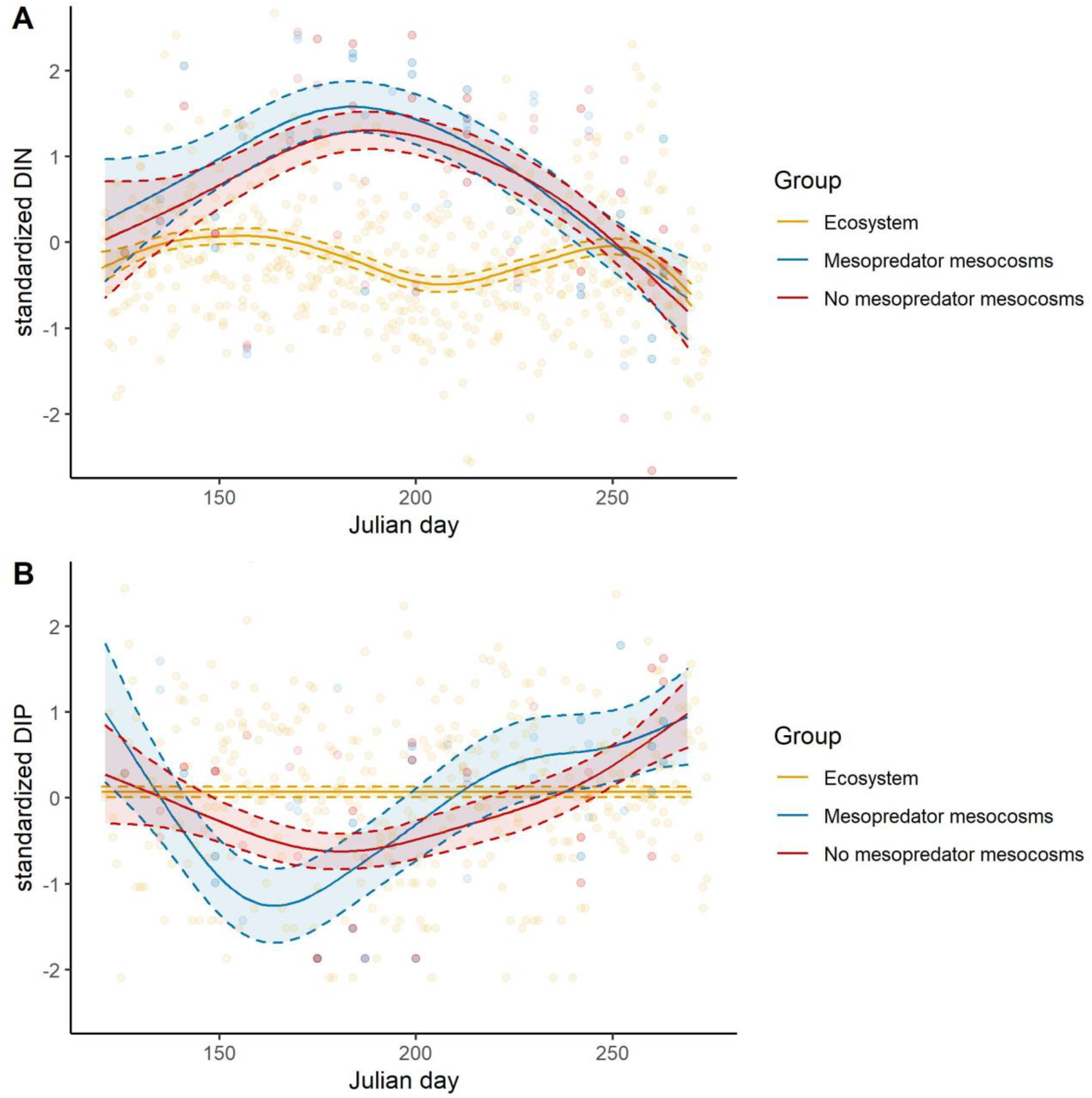
Seasonal development of standardized dissolved inorganic nitrogen (Panel A, sum of nitrate, nitrite, and ammonium = DIN [µml L^-1^]), and standardized dissolved inorganic phosphorus (Panel B, DIP [µml L^-1^]) in mesopredator (blue) and no-mesopredator (green) mesocosm and the ecosystem (orange) for the days 120 – 275 of the years 2015 to 2017. Data points represent independent measurements and were not averaged. The development of both nutrient fractions (points) was modelled (generalized additive model) depending on day of the year (April – October, x-axis) with all replicates being pooled. The solid line represents the mean and the dotted line the 95 % confidence interval. Values of mesocosms and the ecosystem were log-transformed and standardized (z-score), to enable a direct comparison of the ecosystem with mesocosms.

**Table 1.** Median values of monitored parameters in the ecosystem and the mesocosms. Range of values (minimum – maximum) and number of observations is given in parentheses. DIN – dissolved inorganic nitrogen, sum of nitrate, nitrite and ammonium; DIP – dissolved inorganic phosphorus; TN – total nitrogen; TP – total phosphorus; Chl *a* – Chlorophyll *a*; Turbidity – measured at 720nm; Phytoplankton – sum of biovolume per sampling date, only available for end of experiments; Zooplankton – biomass based on geometric conversion (Heerkloss et al. 1991).

|  | Ecosystem | Mesopredator<br>mesocosms | No-mesopredator<br>mesocosms |
| --- | --- | --- | --- |
| DIN ($\mu\text{mol L}^{-1}$) | 1.9 (0.4 – 10, n = 297) | 3.6 (1.2 – 29, n = 47) | 4.2 (0.6 – 12, n = 47) |
| DIP ($\mu\text{mol L}^{-1}$) | 0.06 (0 – 1.8, n = 69) | 0.06 (0 – 0.6, n = 47) | 0.05 (0 – 0.6, n = 47) |
| TN ($\mu\text{mol L}^{-1}$) | 146 (30 – 305, n = 85) | 152 (87 – 285, n = 18) | 230 (95 – 447, n = 18) |
| TP ($\mu\text{mol L}^{-1}$) | 3.7 (1.8 – 7.1, n = 82) | 3 (1.1 – 10, n = 33) | 3.3 (1.2 – 14, n = 34) |
| Chl *a* ($\mu\text{g L}^{-1}$) | 58 (16 – 141, n = 348) | 23 (5.1 – 80, n = 46) | 37 (7 – 300, n = 46) |
| Turbidity ($\text{m}^{-1}$) | 4 (1.1 – 10.1, n = 202) | 2.4 (0.5 – 4.9, n = 43) | 3.1 (1.4 – 16, n = 39) |
| Phytoplankton<br>($\text{mm}^3 \text{L}^{-1}$) | 33 (9 – 83, n = 16) | 17 (4 – 30, n = 4) | 76 (60 – 93, n = 4) |
| Zooplankton<br>($\mu\text{g fresh mass L}^{-1}$) | 754 (42 – 4348, n = 131) | 325 (9 – 9600, n = 37) | 223 (15 – 2800, n = 37) |

Total nitrogen (TN) concentrations within mesocosms developed differently in both treatments already from the start. TN concentrations in mesopredator mesocosms remained around average. TN concentrations in no-mesopredator mesocosms increased throughout the observational period in all years compared to start and end of the experiments as well as the ecosystem (Figure 2A). This development contrasts the DIN development and indicates free DIN was incorporated into other fractions instead. Ecosystem concentrations would decline from June onwards. The TN smooths of the mesopredator treatment was not significantly different to the ecosystem (p = 0.17), but for the no-mesopredator mesocosms (p < 0.0001). Interestingly, the difference in TN between both treatments was not significant (p = 0.22). The trends in both mesocosm treatments points to a higher (no-mesopredator mesocosms), or stable (mesopredator mesocosms) availability of N compared to the ecosystem. Comparable to TN, TP in both mesocosm treatments increased steadily, but mesopredator mesocosms reached average concentrations only at the end compared to no-mesopredator mesocosms (Figure 2B). Ecosystem concentration of TP would start to decrease in July until the end of the observational period. No-mesopredator mesocosm showed an increase of TP starting in July, which indicates a steady P accumulation and is consistent with the DIP increase starting in August. The TP smooths of both treatments were significantly different to the ecosystem for mesopredator (p = 0.03) and no-mesopredator mesocosms (p = 0.0001), as well as both mesocosm treatments to each other (p = 0.03). Mesopredator-mesocosm showed no TP trend from start to end of the experiments, whereas DIP increased from August until the end of September. The DIP:TP remained around 5 – 10% in all mesocosms and the ecosystem (Supplement Figure 3A). The DIN:DIP ratio dropped after August in mesocosms, indicating that DIP was maybe growth limiting during May to June (Supplement Figure 3B). The ratio of TN:TP was highest in mesopredator mesocosms (on average up to 120:1 TN:TP) during July, compared to no-mesopredator mesocosms (on average up to 90:1 TN:TP). Interestingly, the ecosystem showed lowest TN:TP ratios during summer (50:1 TN:TP, Supplement Figure 3C).

**Figure 2.**
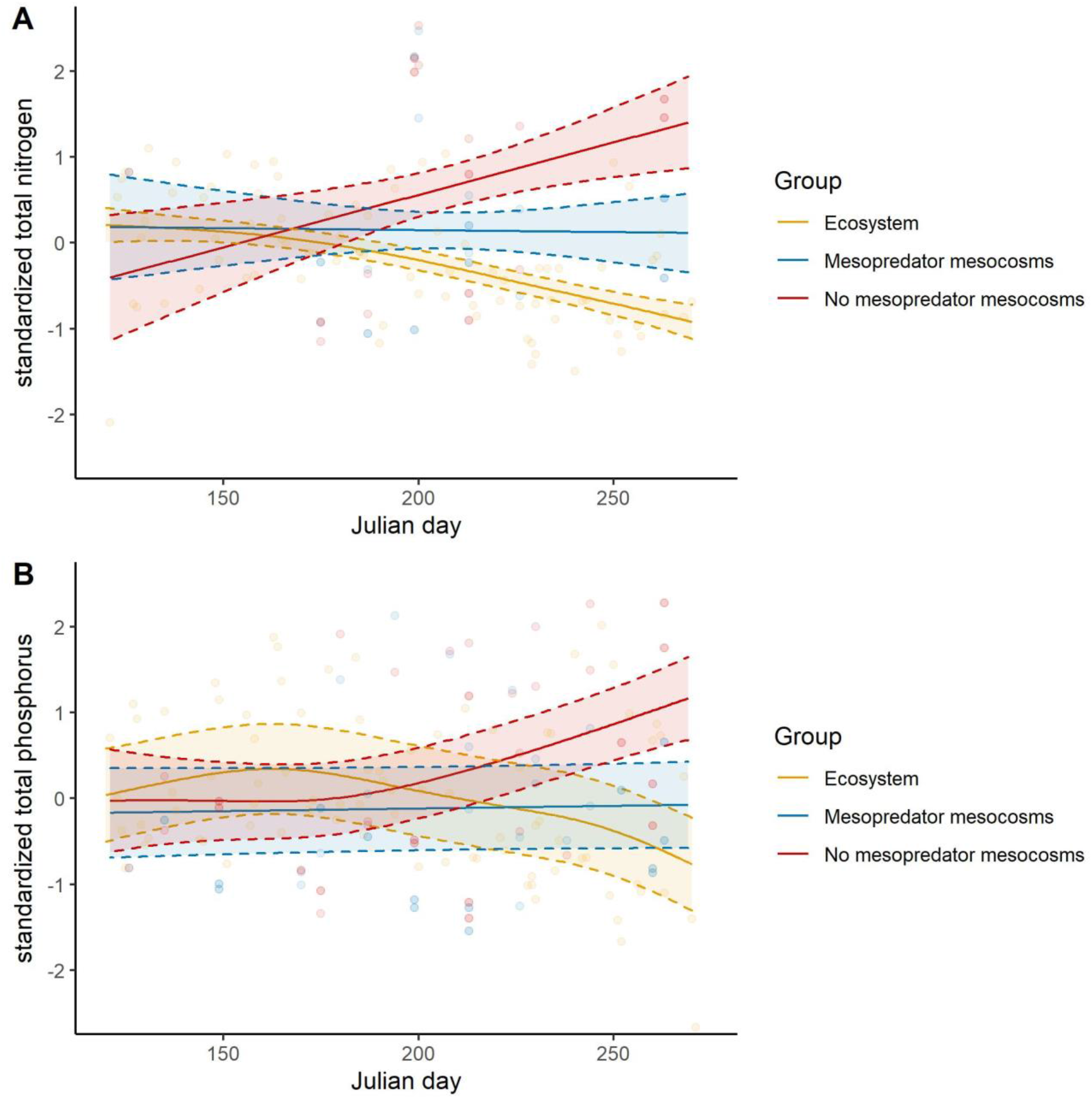
Seasonal development of standardized total nitrogen (Panel A, total N [µml L^-1^]), and standardized total phosphorus (Panel B, total P [µml L^-1^]) in mesopredator (blue) and no-mesopredator (red) mesocosm and the ecosystem (orange) for the days 120 – 275 of the years 2015 to 2017. Data points represent independent measurements and were not averaged. The development of both nutrient fractions (points) was modelled (generalized additive model) depending on day of the year (April – October, x-axis) with all replicates being pooled. The solid line represents the mean and the dotted line the 95 % confidence interval. Values of mesocosms and the ecosystem were log-transformed and standardized (z-score), to enable a direct comparison of the ecosystem with mesocosms.

### Development of planktonic compartments

Chl *a* concentration decreased in both mesocosm treatments below ecosystem average from April until June, when both treatments started to develop independently from each other (Figure 3A). Mesopredator mesocosms remained below average towards the end. Contrary, no-mesopredator mesocosms showed a strong increase of Chl *a* from July until the end of the experiment in September. This pattern coincides with steady nutrient increases in no-mesopredator mesocosms (Figure 3). The ecosystem showed decreasing Chl *a* concentration after the initial spring bloom in April until July. There was a possible small increase of Chl *a* in August suggesting a bloom, which decreased again until the end of September. The Chl *a* smooths of both treatments were significantly different to the ecosystem for mesopredator (p = 0.016) and no-mesopredator mesocosms (p < 0.0001), as well as both mesocosm treatments to each other (p < 0.0001).

**Figure 3.**
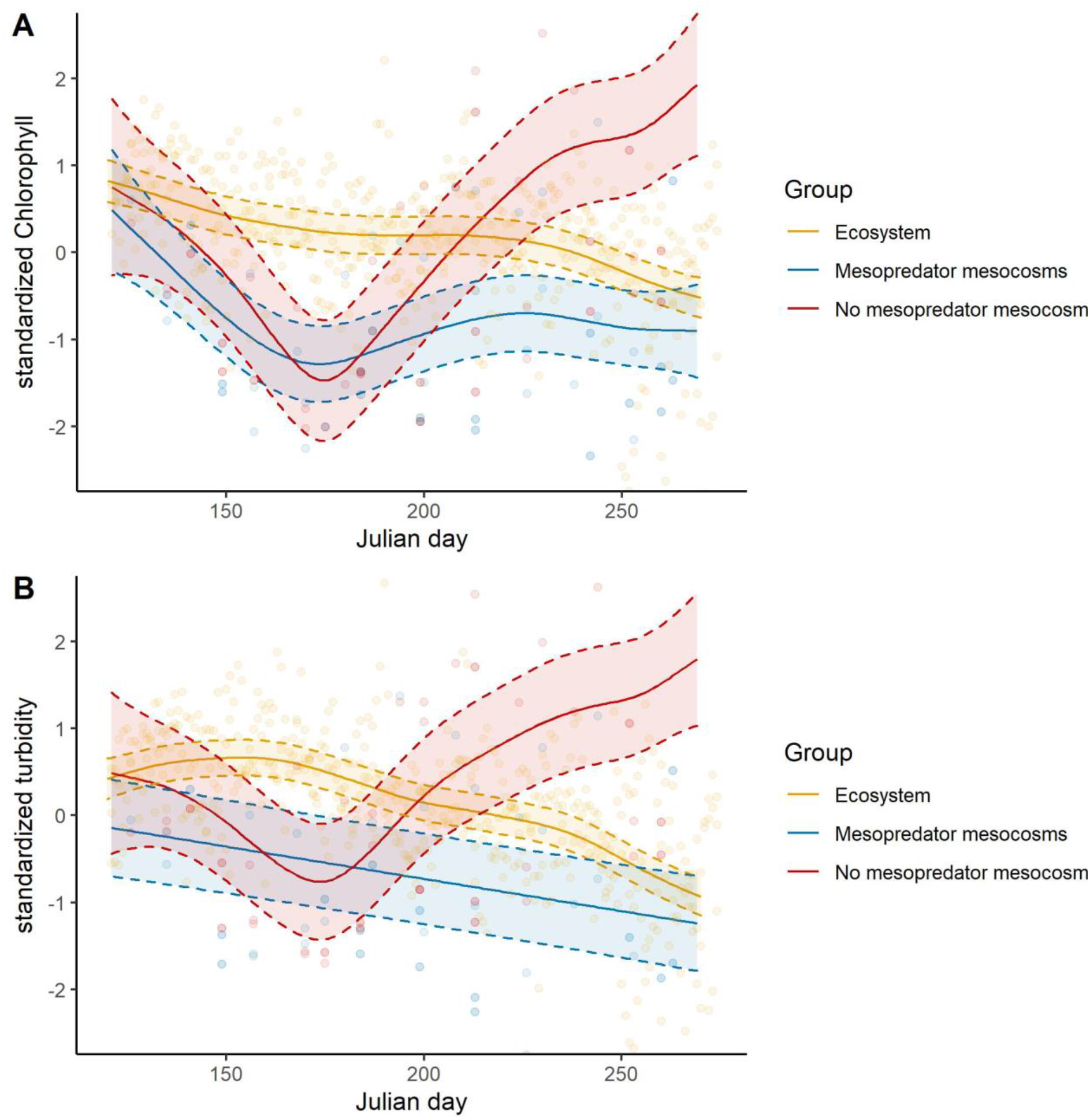
Seasonal development of standardized Chlorophyll *a* (Panel A, Chl *a* [µg L^-1^], proxy for phytoplankton biomass), and standardized turbidity (Panel B, absorption at 720 nm [m^-1^], proxy for light attenuation) in mesopredator (blue) and no-mesopredator (red) mesocosm and the ecosystem (orange) for the days 120 – 275 of the years 2015 to 2017. Data points represent independent measurements and were not averaged. The development of both particulate fractions (points) was modelled (generalized additive model) depending on day of the year (April – October, x-axis) with all replicates being pooled. The solid line represents the mean and the dotted line the 95 % confidence interval. Values of mesocosms and the ecosystem were log-transformed and standardized (z-score), to enable a direct comparison of the ecosystem with mesocosms.

Phytoplankton community composition in the ecosystem was dominated by cyanobacteria across cell volume classes 0.5 – 100 µm^3^ at the start of each experiment (see Supplement Figure 4A). Cyanobacteria were dominated by *Limnothrix* spp., *Synechococcus* spp., and *Aphanothece* spp.. There was an increasing biovolume of larger Chlorophyta (mostly *Scenedesmus* spp.), but median biovolume values were almost one order of magnitude smaller compared to cyanobacteria. Biovolume per size classes changed only slightly in the ecosystem until the end of the experiment (see Supplement Figure 4B). Interestingly, mesopredator mesocosms showed a slightly different phytoplankton community composition with small cyanobacteria still dominating but at a lower biovolume compared to the start of the experiment. Furthermore, other phyla occurred in larger quantities and across size classes, most notably dinoflagellates (for example *Gymnodinium* spp.) and heterokontophytes (*Stephanodiscus* spp.). No-mesopredator mesocosms showed the same phytoplankton community composition as the ecosystem at the end of the experiments but with higher total biovolume. The increased biovolume in no-mesopredator mesocosms corresponds with the higher Chl *a* concentrations also found in those treatments. Heterotrophic bacterial cell count varied widely from 1 to 24·10^6^ cells mL^-1^ and corresponded with overall phytoplankton biomass, that means more phytoplankton, more heterotrophic bacteria. Mesopredator mesocosms showed 30% less bacterial cells mL^-1^ in the available years 2016 and 2017 years compared to no-mesopredator mesocosms. However, a seasonal bacterial development is missing, as counts were only conducted at the end of each experiment.

Likewise, turbidity within both mesocosm treatments decreased initially, but split again after June (Figure 3B). Mesopredator mesocosms continued to show below average turbidity. No-mesopredator mesocosms increased their turbidity in a similar pattern as with Chl *a*. It was not possible to discriminate with the current data set if Chl *a* increased due to increased amounts of particles (abiotic and biotic), or if turbidity increased due to increased Chl *a* (attenuating phytoplankton biomass). Ecosystem turbidity was highest during May and steadily declined afterwards until it was comparable to mesopredator mesocosms turbidity in September. The turbidity smooths of both treatments were significantly different to the ecosystem for mesopredator (p = 0.03) and no-mesopredator mesocosms (p < 0.0001), as well as both mesocosm treatments to each other (p < 0.0001).

Zooplankton biomass (as µg FM L^-1^) per zooplankton class showed different developments within both mesocosm treatments and compared to the ecosystem (Figure 4). Cladocera biomass remained above average in both mesocosm treatments. However, cladocera (*Bosmina* and *Alona* spp.) were rarely detected in the ecosystem, indicating inconclusive trends. The cladocera smooths of both treatments were thus not significantly different to the ecosystem for mesopredator (p = 0.43) and no-mesopredator mesocosms (p = 0.34), as well as not significantly different to each other (p = 0.12). Copepod biomass in mesopredator mesocosms declined steadily after May. Interestingly, the decline of copepod biomass from July on coincides with an increase of Monogononta biomass. Monogononta biomass was average from May until June, reaching above-average biomass during the end of the experiment, contrary to the ecosystem and no-mesopredator mesocosms. No-mesopredator mesocosms showed overall average copepod biomass during the first part of the experiment and a comparable peak like in the ecosystem around in July and August. The copepod smooths of both treatments were significantly different to the ecosystem for mesopredator (p < 0.0001) and no-mesopredator mesocosms (p < 0.0001). However, copepod smooths of both mesocosm treatments were not significantly different to each other (p = 0.26). Monogononta biomass in no-mesopredator mesocosms showed a negative trend with the minima reached at the same time as copepod biomass reached a peak in July and August. Contrary, mesopredator mesocosms showed initially average monogononta biomass, but showed a slight positive trend until the experiment. The monogononta smooths of both treatments were significantly different to the ecosystem for mesopredator (p = 0.001) and no-mesopredator mesocosms (p = 0.0005). Surprisingly, there were no significant differences between both mesocosm treatments to each other (p = 0.24). Differences between mesocosm smooths became more significant for all zooplankton groups, if shorter observational periods were tested, for example from day 150 – day 270 only. However, this also lowered the amount of total observational data. Therefore, we only report this trend but did not analyze it thoroughly. There were no detectable differences in zooplankton species composition between ecosystem and the mesocosms. Cladocera biomass was dominated by *Bosmina* and *Alona* spp., copepod biomass by *Eurytomora affinis* and *Acartia tonsa*, and monogononta biomass by *Keratella* spp., *Filinia* spp., *Brachionus* spp., and *Synachaeta* of various size classes.

**Figure 4.**
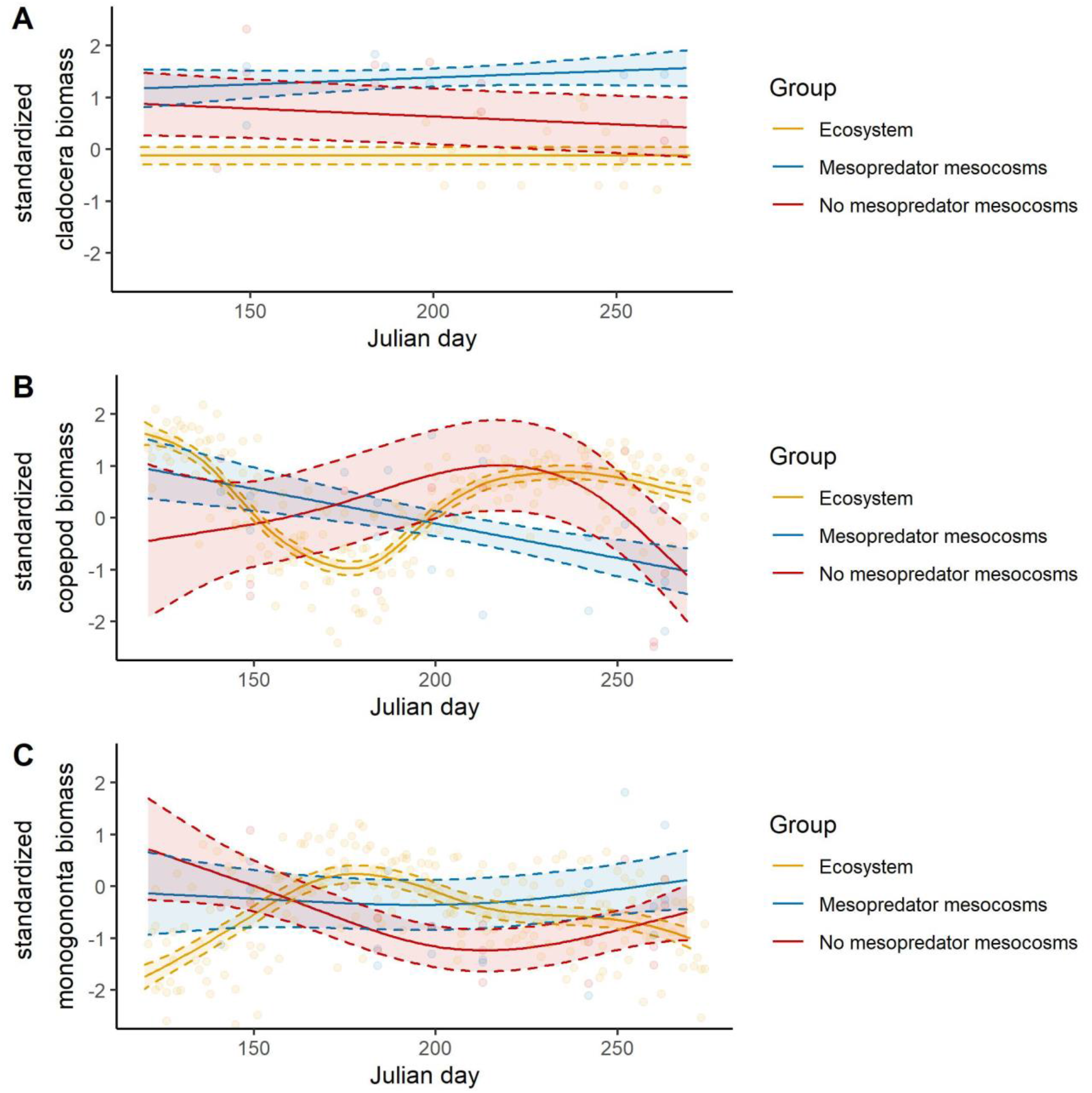
Seasonal development of standardized cladocera (Panel A), standardized copepods (Panel B), and standardized monogononta biomass (Panel C) as fresh mass (µg L^-1^) in mesopredator (blue) and no-mesopredator (red) mesocosm for the days 120 – 275 of the years 2015 to 2017 and the ecosystem (orange, days 120 – 275 of the years 2011 – 2013). Data points represent independent measurements and were not averaged. The development of all zooplankton fractions (points) was modelled (generalized additive model) depending on day of the year (April – October, x-axis) with all replicates being pooled. The solid line represents the mean and the dotted line the 95 % confidence interval. Values of mesocosms and the ecosystem were log-transformed and standardized (z-score), to enable a direct comparison of the ecosystem with mesocosms.

Overall, these results point to an increased grazing pressure on copepods in mesopredator mesocosms, whereas this pressure was less pronounced in no-mesopredator mesocosms, compared to the ecosystem. Interestingly, in mesopredator mesocosms Monogononta biomass was above average in all three years which may indicate a treatment effect by mesopredator occurrence.

Zooplankton within the ecosystem showed again different dynamics. Cladocera density showed no trend, caused by a low number of overall observations, and total biomass. Copepods within the ecosystem declined after high initial numbers in April and showed a second peak later in the year. This development was contrary to both mesocosms, where copepod biomass was average or below average in the later months. Monogononta biomass was highest in June/July, but declined afterwards and remained then lower compared to biomass in mesopredator mesocosms. In conclusion, both treatments had a different impact on abiotic (nutrients) parameters, phytoplankton (Chl *a*), zooplankton, macrophyte and animal biomass (Supplement Figure 5), compared to each other, but also to the ecosystem. These differences are indicative of altered regulating mechanisms caused by the biomanipulation.

### Development of macrophytes and higher trophic levels in mesocosms

Mesopredator mesocosms showed on median more macrophyte biomass compared to no-mesopredator mesocosms, but macrophyte biomass occurred only in three out of five mesocosm replicates for both treatments (Supplement Figure 5). Macrophyte biomass was constituted out of *Charales* (*Chara* sp.) and *Alismatales* (most abundant *Stuckenia pectinata*) in both mesocosm treatments. Mesopredator mesocosms showed more frequent epiphytic growth on mesocosm walls, however this biomass fraction is not represented here, as it was not possible to capture growth and turn-over of this biomass compartment.

Mesopredator mesocosms showed either stable or slightly lower total biomasses of gobies at the end compared to the start of the experiments. However, individual biomass was always increased, pointing to fewer but larger adult animals. Contrary, shrimp abundance increased in four out of five times, with lower individual mass at the end of the experiment, indicating frequent offspring. The one invasive goby *Neogobius melanostomus* found in one mesopredator mesocosm in 2016 had a total biomass of 26 g, and presumably grazed upon the other available gobies, as only two others were found at the end of the experiment in this mesocosm. No-mesopredator mesocosms showed in four out of five replicates colonization with the invasive *Gammarus tigrinus*. Gammarid biomass reached levels of up to 9 g FM m^-2^, whereas only some individuals were ever found in mesopredator mesocosms. This striking difference can be seen in the final animal biomass of both mesocosm treatments (Supplement Figure 5B). Gammarid biomass was so high that total animal biomass was on median as high in no-mesopredator mesocosms as in mesopredator mesocosms. Interestingly, no-mesopredator mesocosms with high gammarid biomass, were always low on epiphytic wall growth, and macrophyte biomass (see Supplement Picture 1 and Supplement Figure 5A). Exemplary gut analyses of gobies showed that they grazed primarily on zooplankton (Supplement Picture 2). Interestingly, an abundance of larger cladocera (*Bosmina* sp. and *Alona* sp.) were found in fish guts, contrarily to the low abundances found in the water column. Exemplary gut analyses of shrimps and gammarids showed mostly macro- and epiphytic biomass.

## Discussion

### Food web impact of biomanipulation

The results of this mesocosm approach needs to be discussed regarding its number of available trophic levels. The no-mesopredator mesocosms showed in all replicates a large biomass of gammarids (representative individuals identified as invasive *Gammarus tigrinus*), with similar or even higher total animal biomass compared to mesopredator mesocosms (Supplement Figure 5). The trophic position of gammarids depends on the presence of grazers and can cause a trophic cascade, that means an influence on other trophic levels, in species-poor ecosystems (Compte et al., 2012). This cascade can result in increased turbidity, macrophyte loss, and increased occurrence of other zooplankton grazers (Compte et al. 2012), similar to our results (see Figure 4). The removal of mesopredator will allow mesograzer to reach a biomass density which is determined by the overall primary production capacity within the ecosystem (Eriksson et al., 2012). However, in the present study, it cannot be concluded that no-mesopredator mesocosms had a higher primary productivity based on higher animal biomass, as gammarids grazed most of the macrophyte biomass, and probably all the epiphyte biomasses (see Figure 5 for the conceptual food web). A recent mesocosm and field study showed that gammarid grazing can cause up to 30% biomass loss of charophytes in the DZLS (Berthold et al., 2022). Such grazing pressure may increase directly with increasing temperatures (Pellan et al., 2016) or indirectly by increased amphipod biomass in a P enriched water column (Östman et al. 2016) probably due to increased food quality of macrophytes (Kraufvelin et al. 2006). A lower macrophyte biomass will cause a decreased possible competition of macrophytes with phytoplankton, and may result in higher Chl *a* values in macrophyte-free mesocosms. A control of these mesograzer by increasing the presence of mesopredator like gobies and shrimp should, therefore, be considered necessary. Gammarids as mesograzer release nutrients by grazing on epiphytes and macrophytes, which can favor phytoplankton. Furthermore, decreased macrophyte biomass lowers direct (nutrients, Kufel & Kufel 2002) and indirect competition (allelopathy, Vanderstukken et al. 2011) with phytoplankton, leading ultimately to a decreased habitat for phyto- and bacterioplankton grazers, like sessile rotifers (Miracle et al. 2007). An increased grazing pressure on gammarids would still allow growth control of epiphytic algae by gastropods (Gagnon et al. 2021), which are very abundant in the form of *Hydrobia* sp. in the coastal waters of the southern Baltic Sea (Paar et al. 2022). Mesograzer like snails can liberate macroalgae from epiphytic growth and recycle necessary nutrients (Bracken et al. 2014). This epiphytic grazing is important especially in turbid coastal waters, where macrophytes are already under light limitation, and epiphytic biomass can increase rapidly during summer months (Paar et al., 2021), possibly causing additional stress on macrophytes (Lin et al., 1996; Özkan et al., 2010).

**Figure 5.**
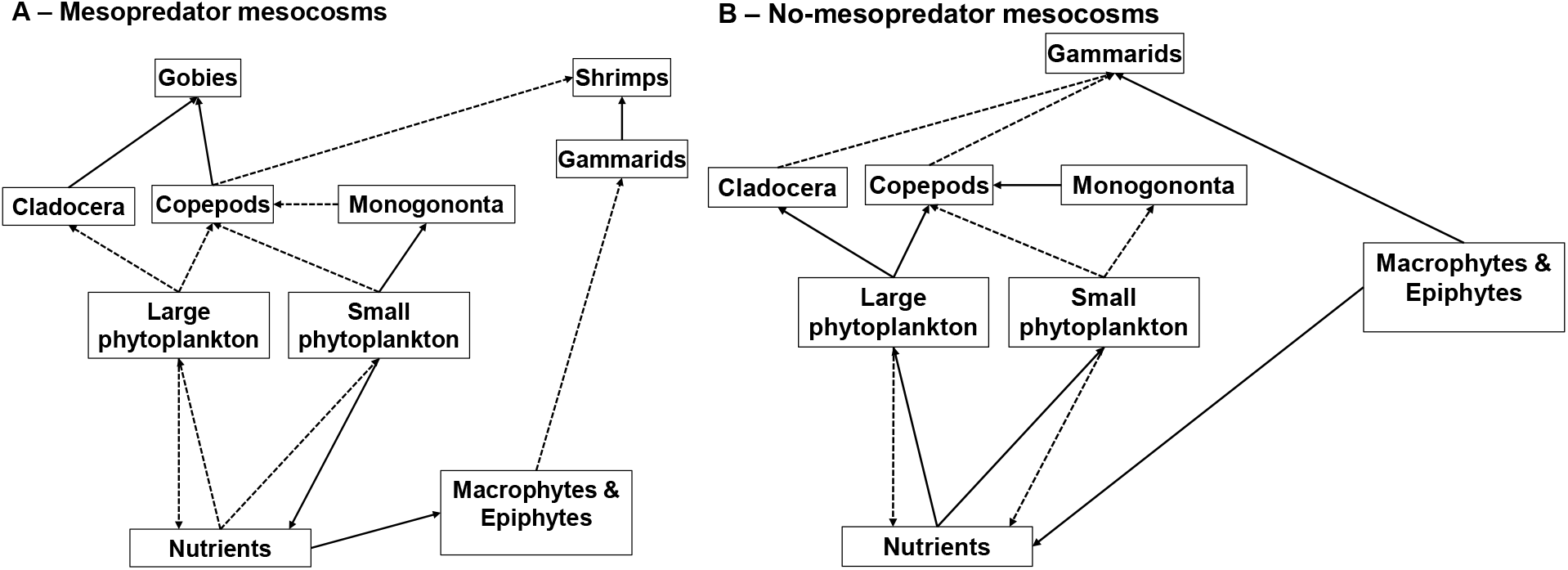
Schematic drawing of the suggested trophic cascade in mesopredator and no-mesopredator mesocosm. Direction of the arrows denotes the direction of matter- and energy flow. Solid arrows denote a strong flow, dotted arrows a weak flow. Bacteria are missing as the potential flows were not established in this study.

The biomanipulation may have impacted nutrient turn-over as well, as pelagic TN and TP stocks increased in no-mesopredator mesocosms (Figure 2), but not in mesopredator mesocosms. This increase was probably mediated through external nutrient inflows by bulk deposition (Supplement Table 1), resulting in lower buffer capacities to external nutrient pressures in no-mesopredator mesocosms. This nutrient increase favored phytoplankton directly, as seen by increasing Chl *a* (Figure 3), as well as probably epiphytic algal biomass (Xie et al. 2013). Ultimately, this increasing phytoplankton biomass will cause high light attenuation which lowers the net-productive part of the water column (Duarte et al. 2002). This increased light attenuation thus results in lower primary production of all compartments (phytoplankton, macrophytes, epiphytes) as seen in lakes (Blindow et al., 2006) and mesocosms (Berthold and Paar 2021).

Mesopredator mesocosms showed lowered turbidity and phytoplankton biomass compared to no-mesopredator mesocosms and the ecosystem. These results are surprising at a first glance, as the occurrence of specific fish species, like three spined stickleback (*Gasterosteus aculeatus*) or cyprinids, are linked to increased turbidity and phytoplankton biomass in mesocosms and ecosystems (Stephen et al., 2004; Williams, Moss, & Eaton, 2002). However, the trophic position as well as function of the occurring fish species needs to be considered. For example, the widely occurring *G. aculeatus* can also cause increasing phytoplankton biomass at abundances of 3 – 6 fish m^-2^ (Jakobsen et al. 2004). This negative effect of *G. aculeatus* may have increased in the Baltic Sea with the decrease of piscivorous fish, like perch and pikeperch stocks (Ojaveer 2002). Surprisingly, grazing on copepods and cladocera in our mesocosms decreased nutrient concentrations, phytoplankton biomass and turbidity nonetheless, which is in contrast to findings in freshwater systems and other lagoons (Feuchtmayr et al., 2009; Jakobsen et al., 2004; Stephen et al., 2004). Thus, *G. aculeatus* and *P. microps/minutus* seem to have different effects on nutrient fluxes within coastal waters. Future studies should analyze if this effect is caused by different feeding behavior and position within the water column, that means bottom dwelling vs. pelagic. For example, a fraction of the nutrients probably went into secondary biomass of mesopredator. Another fraction of nutrients may have been precipitated in the sediments, especially P in the less turbid mesopredator mesocosms (Spears et al. 2008). Our study had a rather large sediment surface and this surface may have been 2 – 3 times larger with infauna (Renz and Forster 2014) thus increasing the effective water-sediment cross-section for nutrient sequestration. There are also other works in freshwater systems, where planktivorous fish had only little effect on grazing control by zooplankton, as phytoplankton grew faster than being grazed upon (Van De Bund et al. 2004, Villena and Romo 2007). Such outcomes can mainly be explained by the additional experimental fertilization, which supports fast-growing or grazing resistant phytoplankton. Especially cyanobacteria can exploit low pulse-wise nutrient supplies (Aubriot and Bonilla 2018) and in conjunction with small cell sizes may be grazing resistant (Lürling 2020). However, environmental daily nutrient depositional rates (see Supplement Table 1) are lower than experimentally added nutrient pulses, and usually do not occur in the form of large, pulse-wise perturbations. Instead, the rate of perturbation is important for organisms and ecosystems to buffer press or pulse disturbances (Shade et al. 2012). In the context of our mesocosms, focusing solely on top-down control allowed to study the impact of altered trophic pathways. The stable TN levels in our mesopredator mesocosms were important for macrophyte establishment, as macrophyte stands tend to decline above thresholds of ∼100 µmol TN L^-1^ (Barker et al. 2008) or under prolonged eutrophic conditions (Coffin et al. 2018). Likewise, a meta-analysis showed that even though fish abundance can increase TP levels, TN is major driver on macrophyte occurrence in already eutrophic lakes (Feuchtmayr et al. 2009). These findings are in line with our results, as TP increased to ecosystem-levels within mesopredator mesocosms but were not causing increased phytoplankton biomasses. Phytoplankton biomass in this part of the lagoon is at least seasonally co-limited for N, as P fertilization experiments in batch tanks and microcosms did not yield any growth response between spring to autumn (Berthold and Schumann 2020). The establishment of macrophytes due to altered nutrient budgets within the water column probably caused several feedback loops. Macrophytes can act as particle traps (Pluntke and Kozerski 2003), however these traits were not observed in these coastal waters (Meyer et al. 2019), or other shallow lakes (Vanderstukken et al. 2011). Nonetheless, submerged macrophytes can improve water chemistry by affecting the sediment microbiome (Ciurli et al. 2009, Zhao et al. 2013), which alters organic C decomposition (Sullivan et al. 2006, Gette-Bouvarot et al. 2015), ultimately lowering O2 respiration in mesopredator mesocosms (Compte et al. 2012). A similar development can also be seen in the areal O2 production of our mesocosms, with mesopredator mesocosms showing a higher net-production with macrophyte establishment in August (see Supplement Figure 1).

Contrary to the mesocosms, the ecosystem showed different dynamics for dissolved nutrients, and decreasing amounts of total nutrients throughout the growth season. The DZLS shows a residence time of 8 – 61 days (Schiewer, 2007), pointing to a steady loss of nutrients after winter nutrient peaks and spring blooms. Macrophytes tend to occur only in shallow regions of the lagoon system, where light is less limiting. However, those local sub-populations are probably especially adapted to low-light climate (Piepho 2017). Therefore, an increase in light penetration depth would allow submerged macrophytes to reach higher biomasses, creating feedback loops like particle trapping, or refuge for herbivorous zooplankton. The question evolves, why phytoplankton biomass remains at a relatively high level within the ecosystem. One cause can be that grazing control on phytoplankton probably decreases, as phytoplankton biomass stabilizes after day 200. This timeframe is in line with the decrease of rotifers biomass (*Keratella* spp.), probably caused by increasing numbers of copepods (*Eurytemora affinis*, Feike & Heerkloss 2009). It is assumed that copepods can cause a grazing pressure on eggs and adult rotifers (Azémar et al. 2007, Feike and Heerkloss 2009). Contrary, the increase in rotifers seems to coincide with the constant below average Chl *a* concentrations and turbidity within mesopredator mesocosms, whereas there is an increase of both Chl *a* and turbidity in no-mesopredator mesocosms. It is therefore possible that grazing pressure within the zooplankton group (intra-guild) caused an inefficient grazing pressure on phytoplankton. These complex trophic niches within zooplankton were recently described for the Baltic Sea, and point to an important impact of rotifers and ciliates on the food web (Zamora-Terol et al. 2020, Novotny et al. 2021).

### Disrupting the microbial loop: A possible switch to treat eutrophic systems?

An increase of rotifer abundance and even diversity was described in top-down manipulated mesocosms (Hansson et al. 2004, Stephen et al. 2004, Miracle et al. 2007), but their impact on pelagic nutrient flows was never clearly established, probably due to simultaneous experimental overload of the nutrient background. Copepod grazing can cause a trophic cascade by grazing on ciliates, which favors heterotrophic nanoflagellates and lowers bacterioplankton. Bacterioplankton is an effective competitor for nutrients with phytoplankton, if organic C is abundant (Joint et al. 2002). The DZLS shows high levels of particulate (16 mg C L^-1^) and dissolved (12 mg C L^-1^) organic C, and bacterial activities (18 µg C h^-1^ L^-1^, Schumann et al. 2003). Thus, efficient grazing on bacterioplankton may favor phytoplankton in otherwise nutrient-limited systems. Likewise, bacterioplankton can also produce extracellular enzymes which indiscriminately support phytoplankton, as nutrient recycling is increased (Chróst and Overbeck 1987). Our limited data suggests that bacterioplankton was lower when mesopredator were present. The number of bacteria corresponded to the total biovolume of phytoplankton cells in the water column, which is another proxy for overall available C in the water column. Thus, a decrease of phytoplankton would cause less bacteria, but the other way around is also possible. However, more data from future studies is necessary to confirm this pattern under these biomanipulative settings. Furthermore, rotifers and ciliates have been described to efficiently graze on small particles and filamentous phytoplankton (Zamora-Terol et al. 2020). Especially filamentous cyanobacteria and picocyanobacteria contribute up to 90% of the phytoplankton biovolume in the DZLS (Albrecht et al. 2017, Berthold and Schumann 2020) and the Baltic Sea (Celepli et al. 2017). Our study found similar phytoplankton community compositions, however, the occurrence of mesopredator increased the absolute biovolume of larger phytoplankton. Indeed, a variety of studies showed that rotifers and other microzooplankton tend to increase with mesopredator presence (Compte et al., 2011; Compte et al., 2012; Šorf et al., 2015). Furthermore, rotifers seem to be more abundant when nutrients are lower, and temperatures increase (Jeppesen et al. 2007, Šorf et al. 2015). This increase in microzooplankton found in our study may have been impacted by a mismatched observational period for zooplankton, as we standardized our mesocosm of 2015 – 2017 with data from 2010 – 2013. However, absolute zooplankton biomass was still higher on median compared to no-mesopredator mesocosms, showing an effect of treatment (see Table 1). The presence of mesopredator which controlled mesograzer and mesozooplankton should have therefore allowed the microzooplankton to increase its abundance, ultimately increasing light penetration depth. This increased light penetration favored macrophytes, which in turn favored even more sessile rotifers among others. A higher amount of macrophytes could potentially allow some biomass to overwinter and create further feedbacks earlier the next year. Contrary, having no grazing pressure on macro- and mesozooplankton did not cause a decrease of phytoplankton biomass, but in conjunction with mesograzer caused a bottom-up control of the ecosystem by releasing more internal nutrients and buffering fewer external nutrients.

### Speculations and Alternative Viewpoints

To apply this food web cascade, there are several possible pathways. For example, an increased fishing quote on large piscivorous fish like perch, pike and sander may increase the abundance of gobies and shrimp, but also sticklebacks and cyprinids. The latter two may increase phytoplankton rather than decreasing it. Large piscivorous fish are already under pressure in German coastal waters and elsewhere in the Baltic Sea. Recreational fishery can catch up to 70% of large piscivorous fish like cod in German coastal waters (Strehlow et al. 2012). Actively fishing for cyprinids and sticklebacks may be an option (Ojaveer 2002), but would simultaneously increase the grazing pressure on gobies and lower the overall food availability for large piscivorous fish. A second option of priming coastal waters with gobies seems more realistic and should be tested in more detail. An increase in goby abundance may create positive feedback loops, like decreasing turbidity. Decreasing turbidity would also improve the sexual selection process of gobies, increasing mating success (Järvenpää and Lindström 2004), thus increasing available food for large piscivorous fish. A combination of management options need to be considered (size and catch limits, time slots) to limit the impact on endangered large piscivorous fish, like cod (Haase et al. 2022), and focus on other non-threatened omnivorous fish.

