## Supplement Data for "Mesopredator-mediated trophic cascade can break persistent phytoplankton blooms in coastal waters"

### Methodological remarks on a possible “bottle effect” within the mesocosms

Every *ex situ* or invasive approach will eventually lead to a deviation from the actual ecosystem. To keep the comparability with the ecosystem high, we used mesocosms with larger volumes and surface areas, representative water depth for the DZLS that still allowed macrophyte growth, and interannual replicates. Furthermore, results of the mesocosms were standardized with ecosystem data to allow a direct comparison. Abiotic factors that can be impacted by *ex situ* approaches are temperature, oxygen, and water levels. Nutrient fluxes into the mesocosms can lead to a buildup of elemental stocks, as there is no water exchange.

Temperature was relatively stable from surface to bottom with differences of ±0.5 °C 95% of the time (Berthold 2018) and was comparable with other larger mesocosms which used active rather than passive cooling (Wahl et al. 2015, Pansch et al. 2016). Surface temperature was only in 3.5% of time > 2 °C compared to bottom temperatures (Supplement Figure 6A) and only in the first weeks of June 2015 until pump positions were improved. Oxygen concentration was on median above 8 mg L^-1^ regardless of surface or bottom measurements (Supplement Figure 6B). Furthermore, the areal O_2_ production (mmol O_2_ m^-2^ d^-1^) was always net-positive in 2015 (see Supplement Figure 8), which contrasts with the net-heterotroph Zingster Strom within the DZLS for the same time (see Supplement Figure 9). However, the different water depth needs to be considered, with the mesocosms being probably not light limited, whereas the Zingster Strom shows a steep light gradient within the first meters (Schubert et al. 2001). Interestingly, the results of this net-autotrophy in larger mesocosms is in contrast to small mesocosms measuring mostly net-heterotrophy even in the presence of abundant phytoplankton or macrophyte biomass in a shallower water column (35 cm instead of 70 cm, Berthold and Paar 2021). One cause can be that the total sediment area to water column is lower in the larger mesocosms, which results in lower relative respiration of sediment in deeper mesocosms compared to shallower mesocosms. The water level was less of a concern, as the mesocosms were open to the atmosphere and water deposition by rain was constant with evaporation. Salinity within the mesocosms ranged on median around 6.8 ± 0.7 PSU. Nonetheless, for some occasions the addition of deionized water during long-term draughts was necessary, as otherwise primary productivity rates of macrophytes would have been affected (Bucak et al. 2012). However, this openness to the atmosphere can also be a problem during periods of extensive bulk deposition of N and P by rain and dust. Precipitation in this area is highest during summer months, as well as areal PO_4_ and total P deposition (µmol m^-2^ d^-1^, Berthold et al. 2019). Out of the monitoring data set of the Biological Station Zingst, we could calculate a net flux of 26 – 38 µmol L^-1^ DIN, 52 – 67 µmol L^-1^ total N, 0.3 – 1 µmol L^-1^ DIP, and 1.9 – 4.7 µmol L^-1^ total P into the mesocosms (see Supplement Table 1 for specifics). Interestingly, these additional fertilizations were not recognizable in the total N or P pool within mesopredator mesocosms. However, no-mesopredator mesocosm were more affected, which may not only point to difference in buffer capacities to external nutrient supply, but maybe also to a different food web mediated nutrient flow.

Supplement Table 1 Depositional rates of nutrients, nitrogen (N) and phosphorus (P) in dissolved (DIN and DIP) and total form (TN, TP) from April to September (2015 – 2017). Total deposits are the sum of all monitored nutrient inputs by bulk (rain and dust) deposition, monitored by the Biological Station Zingst. Missing values compensated by an average share of dissolved to total nutrients (see Berthold et al. 2019). Volumetric increase is the total deposit divided by the mesocosm volume (200 L) to show the probable concentration increase per volume.

| Year | DIN | | DIP | | Total N | | Total P | |
| --- | --- | --- | --- | --- | --- | --- | --- | --- |
|  | total deposit  (mmol) | volumetric increase (µmol L^-1^) | total deposit  (mmol) | volumetric increase (µmol L^-1^) | total deposit  (mmol) | volumetric increase (µmol L^-1^) | total deposit  (mmol) | volumetric increase (µmol L^-1^) |
| 2015 | 75.9 | 38.0 | 1.0 | 0.5 | 122.7 | 61.3 | 8.5 | 4.2 |
| 2016 | 52.4 | 26.2 | 1.9 | 1.0 | 103.4 | 51.7 | 9.4 | 4.7 |
| 2017 | 72.8 | 36.4 | 0.6 | 0.3 | 134.3 | 67.2 | 3.8 | 1.9 |

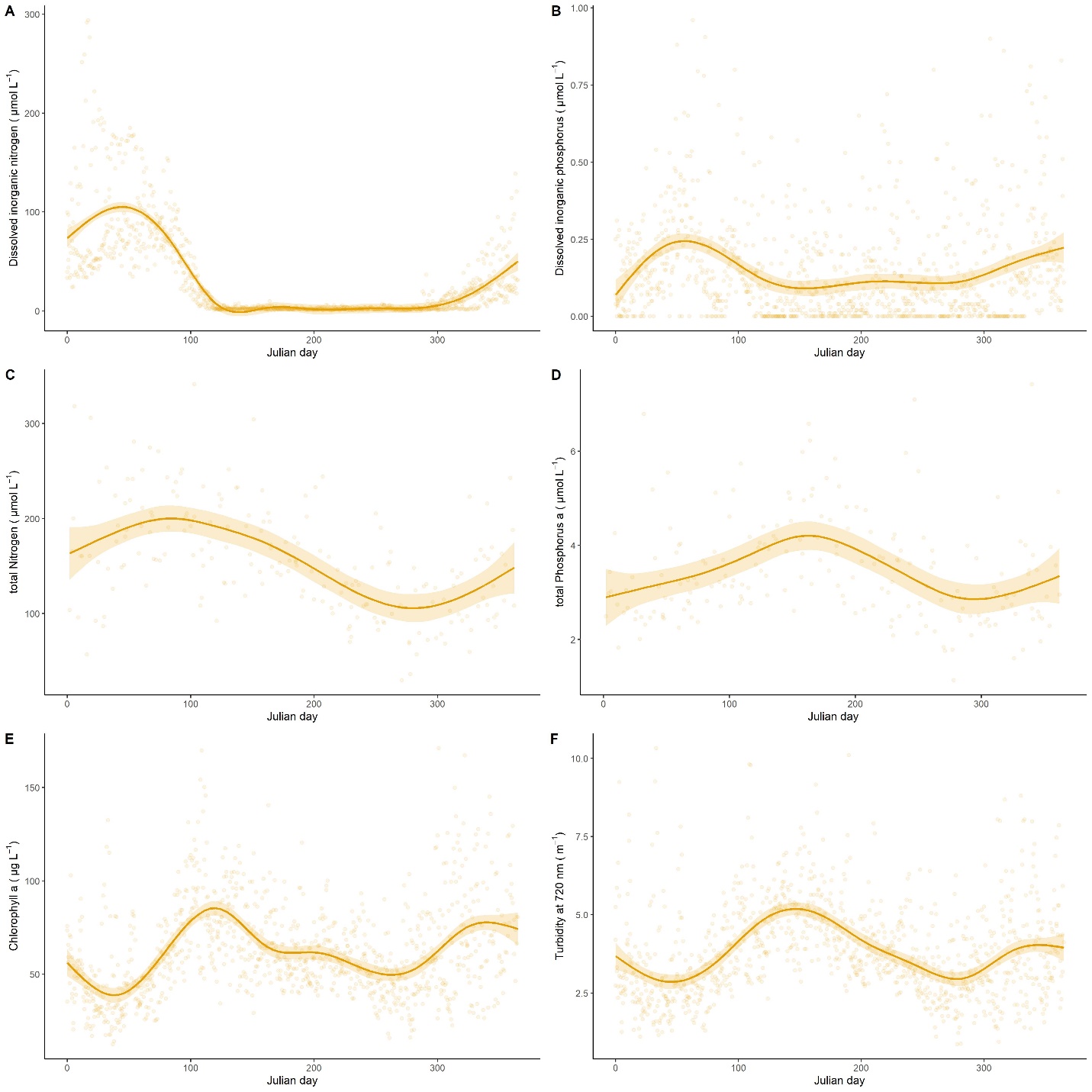

Supplement Figure 1 Annual development (based on Julian day) of abiotic and biotic parameters in the Darss-Zingst lagoon system pooled for the years 2003 – 2017. Dots represent actual data points, the solid line represent the average (smoothed by generalized additive modelling, function geom_smooth, “gam”, Wickham 2016), and the colored ribbons the ±95% standard error.

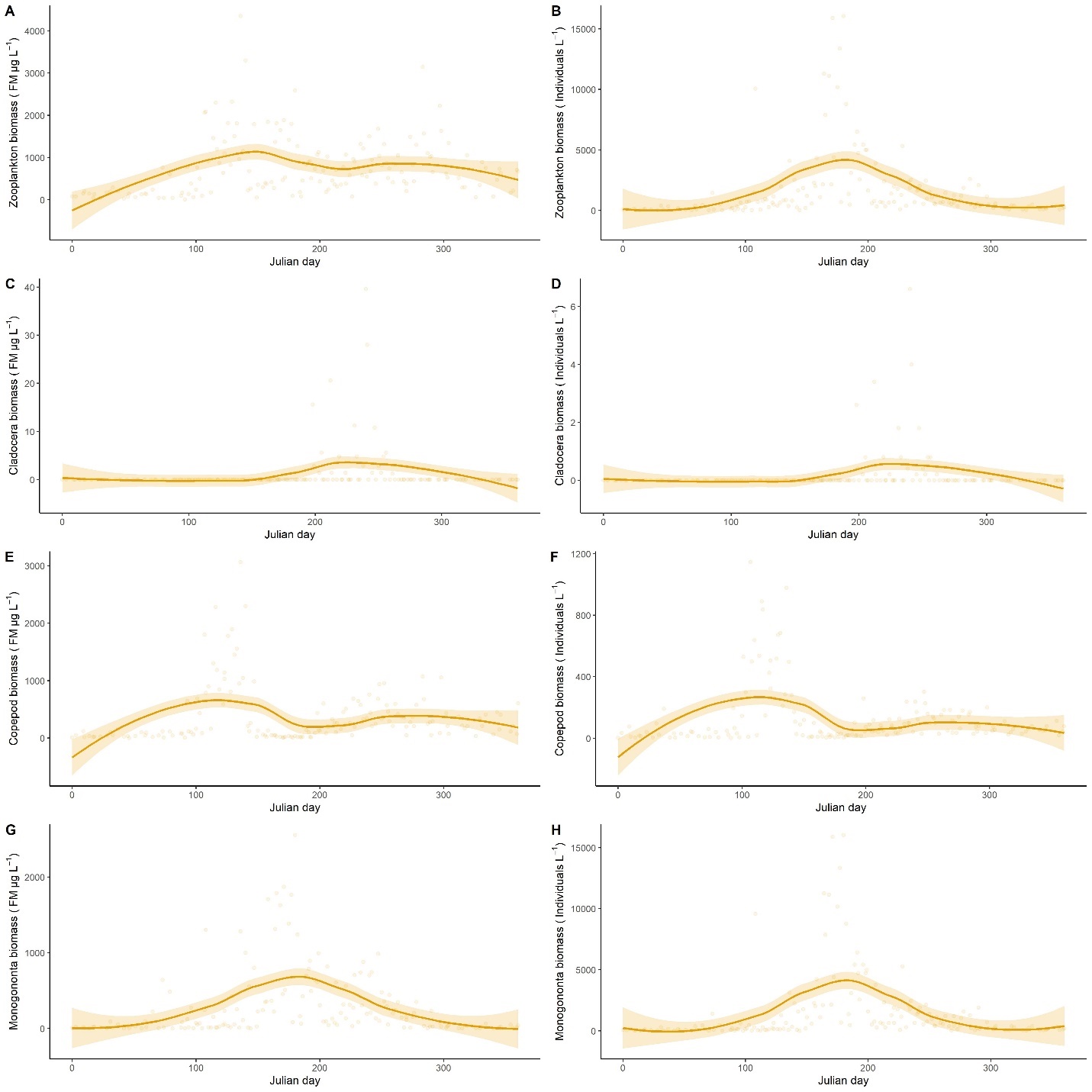

Supplement Figure 2 Annual zooplankton development (based on Julian day) of biomass (fresh mass in µg L^-1^), and density (Individuals L^-1^) of total zooplankton, cladocera, copepod, and monogononta in the Darss-Zingst lagoon system pooled for the years 2003 – 2013. Dots represent actual data points, the solid line represent the average (smoothed by generalized additive modelling, function geom_smooth, “gam”, Wickham 2016), and the colored ribbons the ±95% standard error.

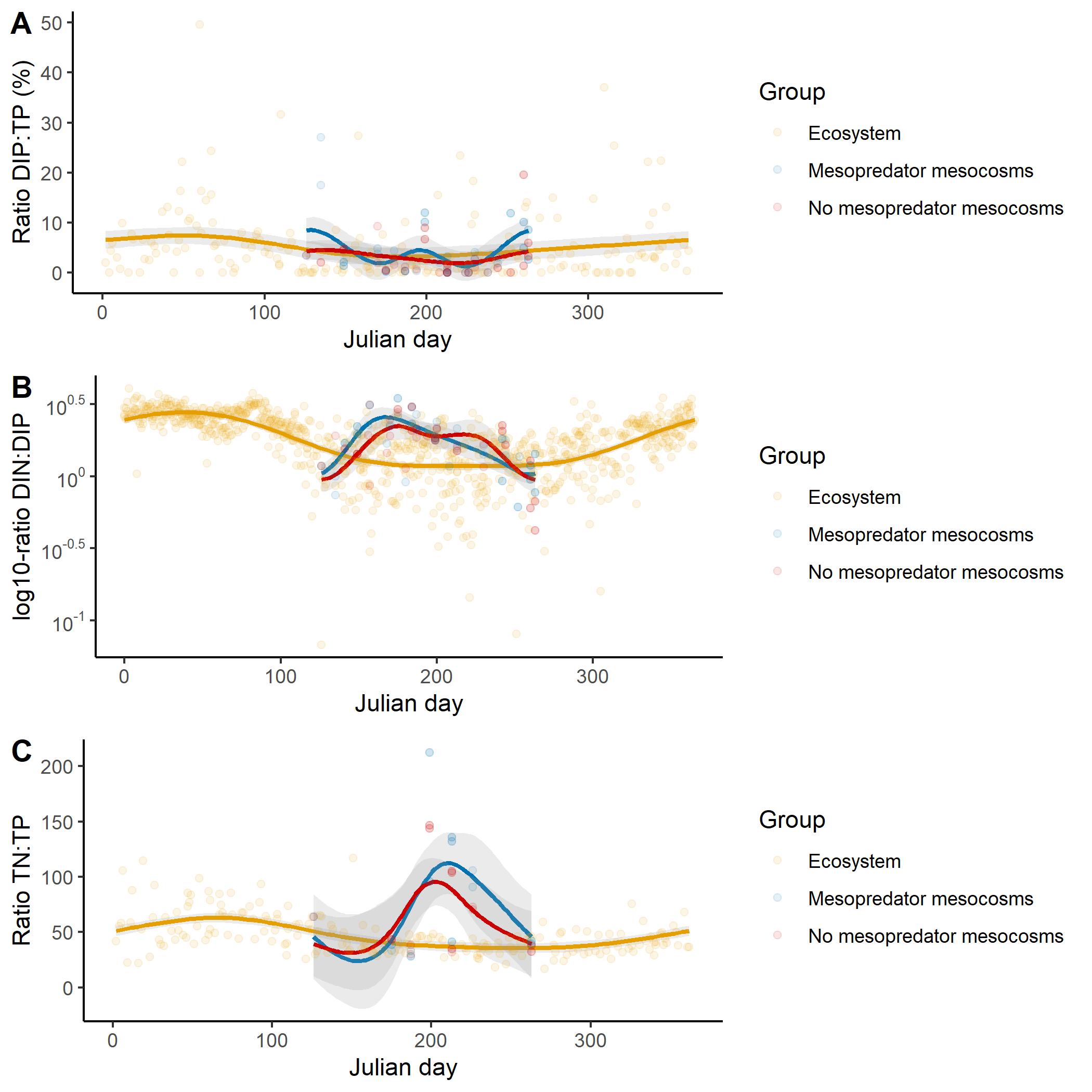

Supplement Figure 3 Annual development (based on Julian day) of various ratios for dissolved phosphorus (DIP) to total phosphorus (TP), dissolved nitrogen (DIN) to DIP, and total nitrogen to TP, in the Darss-Zingst lagoon system (gold) pooled for the years 2003 – 2017 and the fish (blue) and no-mesopredator mesocosms (green). Dots represent actual data points, the solid line represent the average (smoothed by generalized additive modelling, function geom_smooth, “gam”, Wickham 2016), and the colored ribbons the ±95% standard error. Please note the log-scale on B, as DIN:DIP ratios can be as high as 12000:1 during winter conditions (very high DIN, very low DIP).

Supplement Figure 4 Whisker-Box plots of phytoplankton community biovolume (mm^3^ L^-1^) sorted by cell size classes and phytoplankton phylum. Data represents the ecosystem communities at the experimental start (May of each year 2015 – 2017) used for filling the mesocosms (Panel A), and at the end of the experiment (September of each year 2015 – 2017) in the ecosystem, mesopredator and no-mesopredator mesocosms (Panel B). Phytoplankton samples for the ecosystem were based on mean values of at least four independent sampling dates.

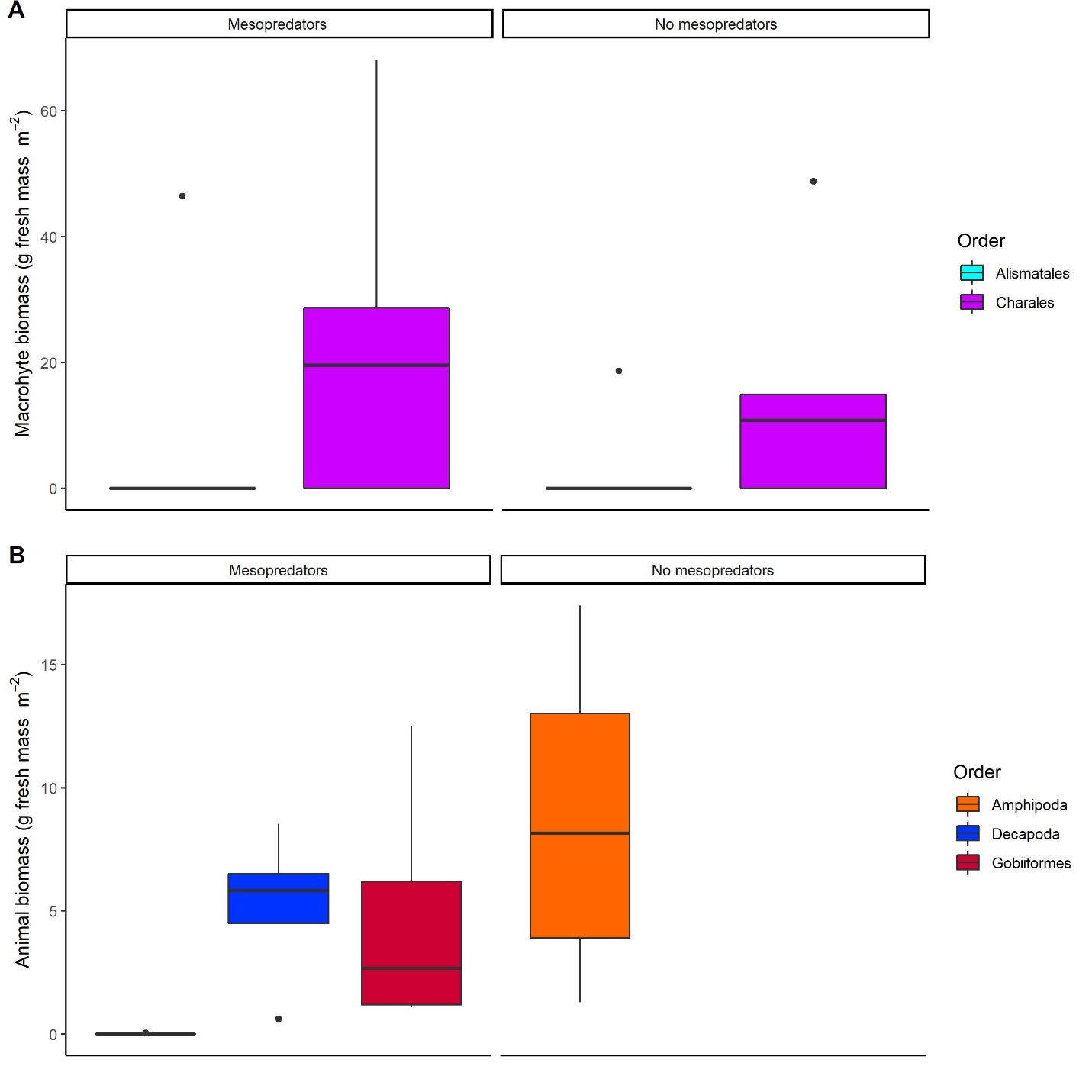

Supplement Figure 5 Whisker-Box plots of macrophyte (Panel A) and animal biomass (Panel B) separated on order and based on g fresh mass per m^-2^ in mesopredator and no-mesopredator mesocosm after the end of each experimental run of the years 2015 to 2017. Macrophyte biomass included epiphytes on the macrophytes, if present.

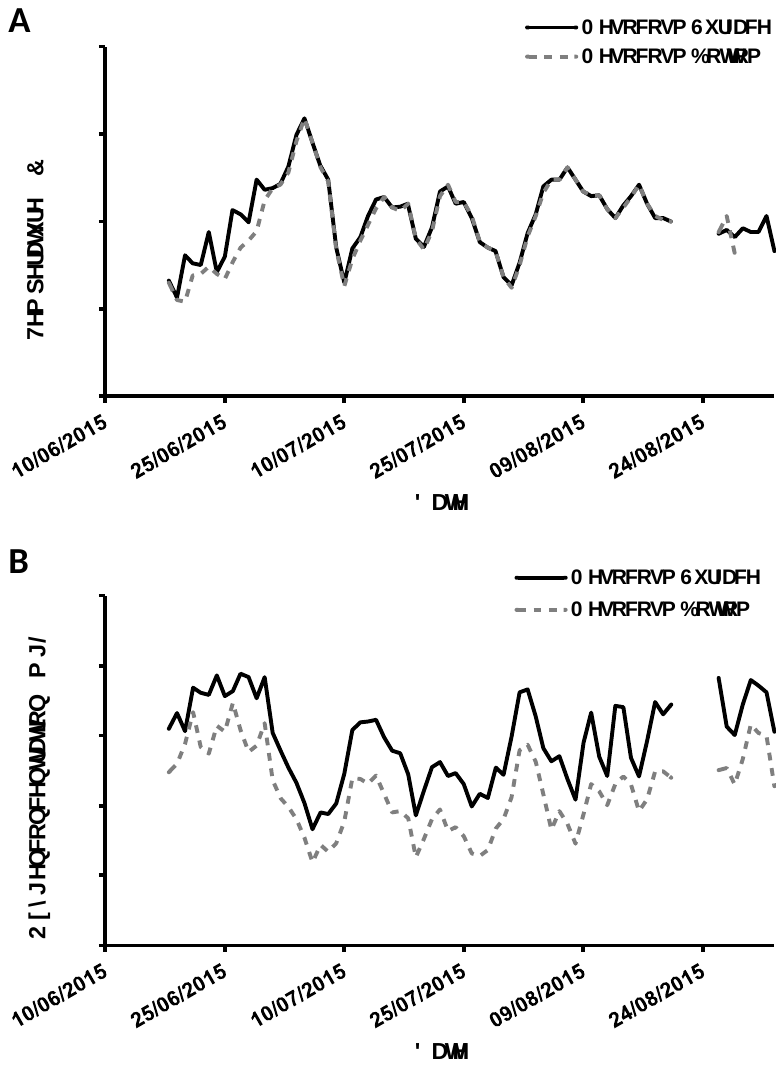

Supplement Figure 6 Daily running median temperature (°C, Panel A) and oxygen (mg L^-1^, Panel B) at the mesocosm surface (5 – 20 cm water depth) and the bottom (70 cm water depth). Temperature and Oxygen was measured every 5 minutes simultaneously at surface and bottom (n = 20588).

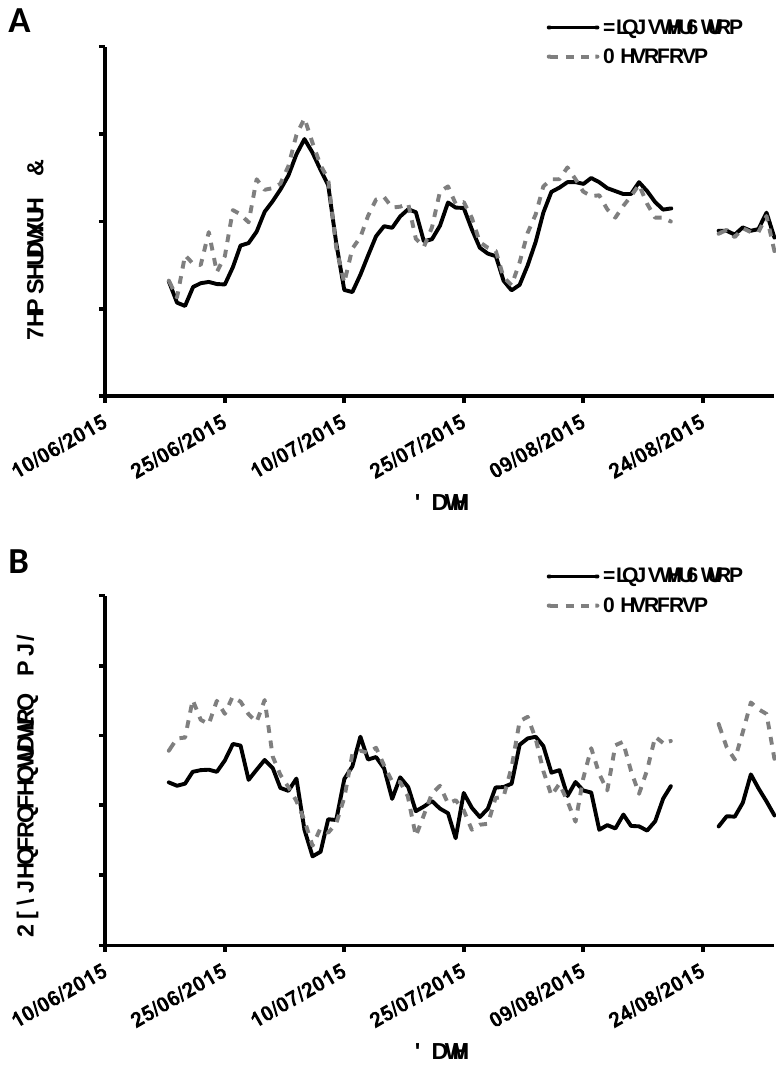

Supplement Figure 7 Daily running median temperature (°C, Panel A) and oxygen (mg L^-1^, Panel B) in the mesocosm (mean value between bottom and surface) and the Zingster Strom (50 cm water depth). Temperature and Oxygen was measured every 15 minutes inside the Zingster Strom (n = 9209) and every 5 minutes in the mesocosms (n = 20588).

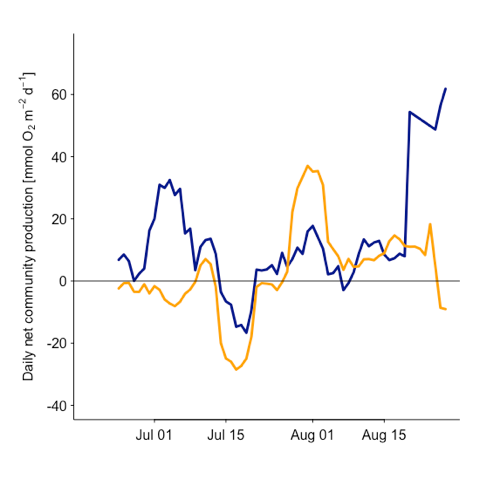

Supplement Figure 8 Depth-integrated daily net community production (mmol O_2_ m^-2^ d^-1^), the sum of daytime production and nighttime respiration in mesopredator mesocosms (blue), and no-mesopredator mesocosms (orange) in 2015. Oxygen saturations were measured every 5 min with optodes at the water surface and recalculated to oxygen concentrations based on temperature and salinity. Daytime respiration was calculated from the average nighttime respiration rate of the respective system and multiplied by the hours of daylight. Dates of cleaning and sampling including the following day were removed prior the analysis. Weekly running mean of daily values are shown.

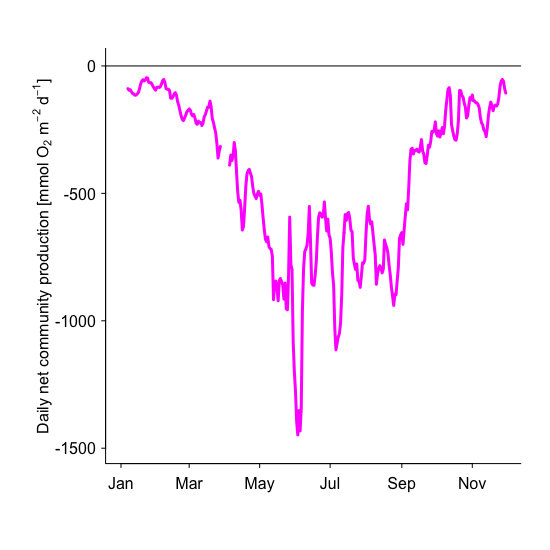

Supplement Figure 9 Depth-integrated daily net community production (mmol O_2_ m^-2^ d^-1^), the sum of daytime production and nighttime respiration in the Zingster Strom in 2015. Oxygen saturations were measured every 15 min with optodes at the water surface and recalculated to oxygen concentrations based on temperature and salinity. Daytime respiration was calculated from the average nighttime respiration rate of the respective system and multiplied by the hours of daylight. Dates of cleaning and sampling including the following day were removed prior the analysis. Weekly running mean of daily values are shown.

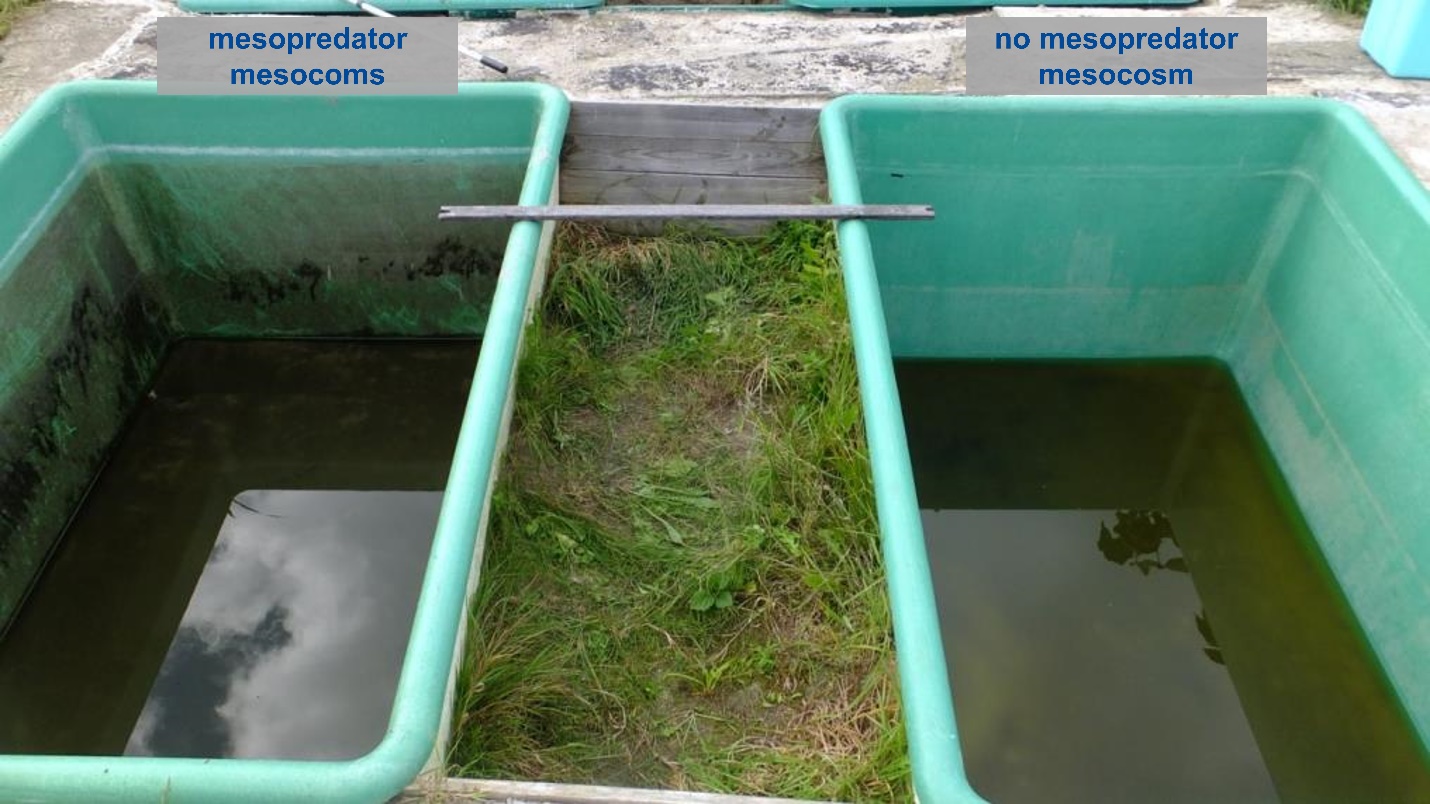

Supplement Picture 1 Photography of two mesocosms at the end of one experimental run on the 18^th^ September 2017, left with mesopredator present, right without mesopredator present.

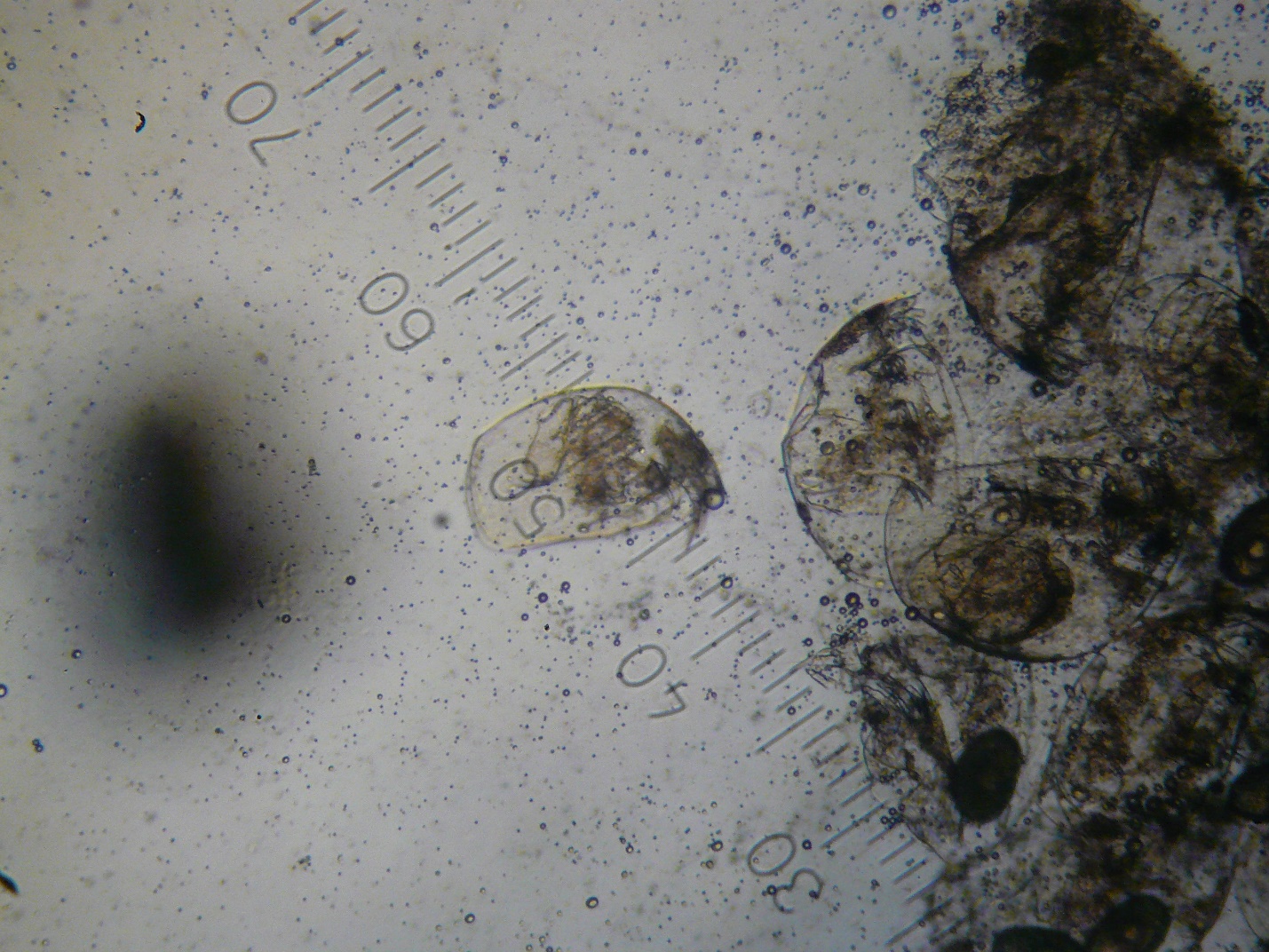

Supplement Picture 2 Exemplary gut analyses of one *Pomatoschistus* *microps* out of a mesopredator mesocosm on the 11^th^ November 2015. Large cladocera were found throughout.
